# Bio-contaminated Plastic Micropipette Tip Sterilization Stations: Environmentally, Economically, and Energetically Viable Solution

**DOI:** 10.1101/2023.12.21.572721

**Authors:** Arian Veyssi, Laxmicharan Samineni, Rashmi P Mohanty

## Abstract

Bioscientific research laboratories significantly contribute to global plastic waste production through their widespread use of plastic products, such as single-use micropipette tips. However, biologically contaminated pipette tips must undergo several washing and sterilization steps before being reused or recycled. Grenova Solutions provides such a decontamination station called TipNovus, which has been implemented by academic and government research labs to reuse pipette tips in sensitive biological assays. Despite this success, the high initial purchasing cost of these washing stations deter many laboratories from incorporating it into their workflow. Additionally, researchers are reluctant to reuse pipette tips due to concerns that the washing process may not thoroughly remove all contaminants. To mitigate these concerns, considering the University of Texas at Austin as an example, we performed a cost-benefit analysis of employing a university-wide washing station. We estimated that only single-time reuse of the pipette tips could result in a 100% return on investment from the equipment purchase cost within 3 months. Then, with our pilot experiments, we confirmed the TipNovus washing steps to be 100% efficient in sterilizing pipette tips contaminated with T7 bacteriophage, enabling their reuse in bacteriophage functionality assays. Finally, we proposed an alternative and more convenient autoclave-based sterilization method to decontaminate pipette tips.

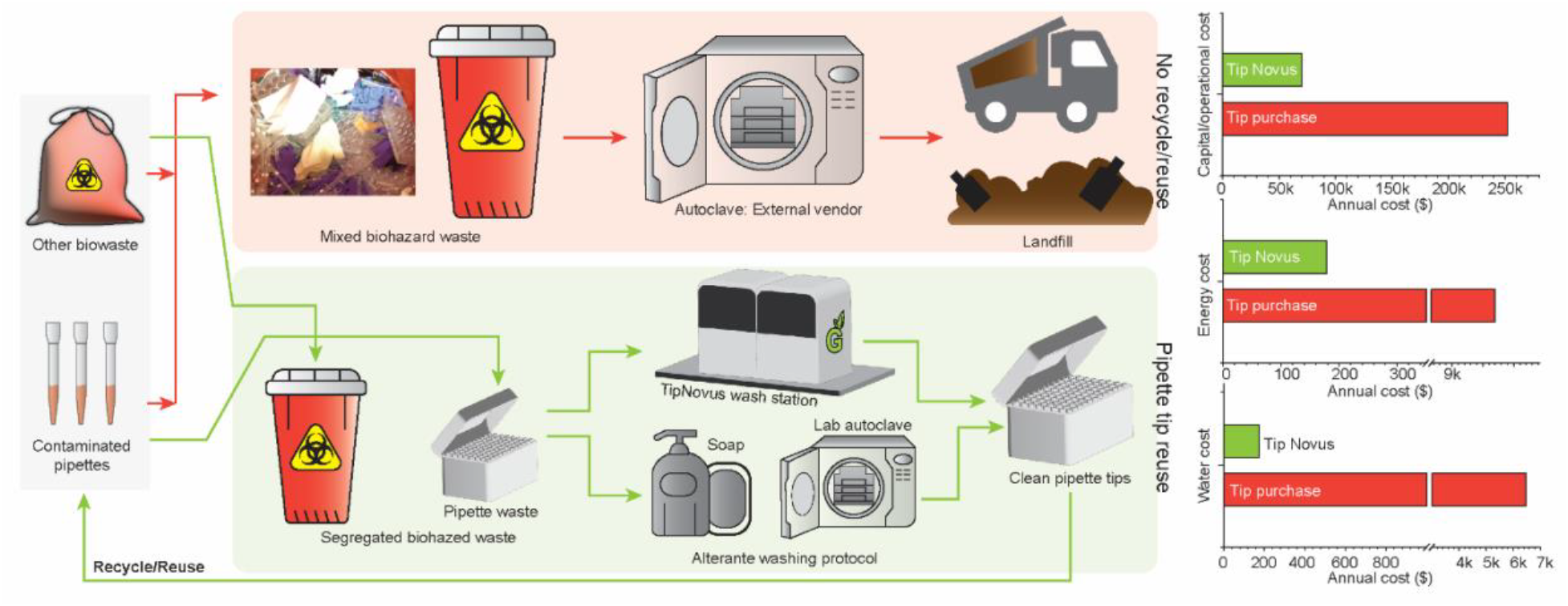

**SYNOPSIS:** Single-use plastic micropipette tips discarded by bioresearch labs generate substantial waste. This study reports adopting standardized tip-washing and reuse systems in labs greatly reduces plastic waste and research expenditures.

## INTRODUCTION

Research laboratories are a major contributor to non-biodegradable plastic waste due to the extensive use of disposable plastic items, including pipette tips, microcentrifuge tubes, and Petri dishes. Around 5.5 million tons of plastic waste were generated from bioscientific research laboratories in 2014 alone, contributing to 1.8% of the total plastic waste produced globally.^1,2^ Recycling this waste would save 1 billion gallons of gasoline and reduce carbon dioxide emissions by 16.5 million tons.^3,4^ Biological research labs, especially, use copious amounts of single-use plastic micropipette tips during day-to-day operations, which costs them millions of dollars. After a single use, the bio-contaminated pipette tips are autoclaved and end up in landfills or incinerators.

While researchers and universities practice sustainability in their everyday lives, they are skeptical about implementing these practices in their research. The research community is constantly investigating innovative strategies such as replacing fossil fuel-derived single-use lab consumables with bioplastics,^5^ and discovering plastic degrading enzymes to limit the environmental plastic burden.^6,7^ Apart from these attempts, it would be beneficial to focus efforts on implementing the three fundamental steps in the plastic waste management hierarchy, the three Rs: Reduce, Reuse, and Recycle.^8,9^ Microbiology research labs that have strived toward sustainable lab practices by incorporating various reduction and reuse approaches have eliminated up to 43 kg of single-use plastic waste in 4 weeks and reduced total lab expenditures by up to 40%.^10,11^ To reduce the waste generated from disposable pipette tips, research labs have implemented the following effective strategies: reusing tip boxes, procuring pipette tip boxes or packages produced from sustainable materials, using smaller pipette tips when appropriate, and employing washable or reusable alternatives.^8,9,11^ Although the purchasing cost of glass and metal alternatives is higher than plastics, they can be washed, sterilized, and reused conveniently, rendering them both environmentally and economically viable.^9,11^

Replacing single-use plastics in biological labs is not always viable, as the chosen material must meet the sterility and non-pyrogenic requirements. When more suitable alternatives to replace plastics do not exist, efforts should be made to reuse them. However, when it comes to reusing plastic in the lab, contamination is the central issue, as laboratory work necessitates high sterility. Plastics that encounter biological contaminants require laborious decontamination procedures before being reused. The next promising solution to mitigate plastic waste is recycling, which reduces environmental impact and greenhouse gas emissions. Recycling biologically contaminated plastic waste also requires extensive washing and decontamination procedures.^4^ Therefore, it is crucial to prioritize the establishment of dedicated washing and decontamination stations to reuse or recycle contaminated plastic pipette tips.

A high throughput, automated pipette tip washing and decontamination station, TipNovus, is commercially available from Grenova.^12^ This washing system has enabled the reuse of 93 million pipette tips with an average economic savings of $112-$168 per hour.^9,12^ TipNovus has been installed by academic research labs, pharmaceutical companies, and government labs to reuse plastic pipette tips in biological experiments, including mass spectroscopy, polymerase chain reaction (PCR), next-generation sequencing (NGS), enzyme-linked immunosorbent assay (ELISA), and RNA library production. Despite this success, most academic research labs are reluctant to incorporate these washing stations into their workflow.^9^ One of the concerns is the high capital cost of installing the equipment in individual research laboratories. Another concern is the effectiveness of the washing process. The details of the tested contaminants and assays used to confirm their absence after a single wash cycle are not readily available. Due to apprehension in the academic community, successful implementation of pipette tip reuse for sensitive and novel biological experiments would require studies to validate the efficiency of washing protocols for various scenarios and establish the economic and environmental impact of implementation.

Here, considering the University of Texas (UT) at Austin as an example, we performed a cost-benefit analysis demonstrating the economic and environmental benefits of installing a university-wide micropipette-tip washing station. Next, we designed a pilot experiment to evaluate the effectiveness of the washing steps implemented in the TipNovus system. Using T7 bacteriophage, a bacterial virus, as one of the biological contaminants, we estimated the percentage of contaminants eliminated from the pipette tip surface after a single washing and sterilization cycle. We then investigated the reusability of the washed pipette tips. Further, we have introduced and verified a more inexpensive autoclave-based pipette tip decontamination method, which is already in place to reuse glassware in biological research laboratories as an alternative to a centralized washing system.

## MATERIALS & METHODS

### Materials and Equipment

T7Select® Packaging Kit (MilliporeSigma), BL21 *E. Coli*. bacterial strain (Novagen), M9LB agar media (Thermo Fischer Scientific), dishwashing detergent (Dawn), UV incubator (Analytik Jena, #UVP SI-950), ultra sonicator (Branson Ultrasonics™, #CPX952338R), 200 µL pipette tip racks (Fisherbrand, #02707417), autoclave (Consolidated Sterilizer Systems, #SR-24B-ADV-PRO), Tris-HCl (Fisher Scientific, #BP153-500), NaCl (Fisher Scientific, #S271-3), MgSO_4_ (Fisher Scientific, #BP213-1)

### Contamination of plastic pipette tips with T7 bacteriophage

To prepare contaminated pipette tips that would be used to validate the washing protocols tested in this study, we rinsed five tips per study group in 2.30 × 10^9^ plaque-forming units (pfu) or 4.80×10^9^ pfu of T7 bacteriophage stock. We inserted them into empty pipette tip racks. Next, we stored one group of the contaminated tips at 4°C until phage quantification and termed them as ‘unwashed tips.’ The other group, ‘washed tips,’ was first autoclaved for 30 minutes of sterilization cycle at 121°C and 100 kPa and then stored at 4°C until further washing steps.

### Washing protocol 1. Pipette tip washing to replicate TipNovus sterilization

To validate the washing procedure employed by TipNovus, we replicated the process in the lab based on conversations with Grenova. From our conversation with Grenova, we found that TipNovus washing comprises a 3-stage washing and sterilization process: washing, sonication, and Ultraviolet C (UV-C) sterilization.^12^ While a high-pressure washing step eliminates large contaminants from the pipette tip surface, ultrasonic cleaning removes small contaminants and particles. The final UV sterilization kills infectious contaminants from the surface. We replicated this protocol by sequentially implementing each step of the washing process. In step 1, the high-pressure washing step was reproduced by first soaking the autoclaved tips in high-quality detergent for 1 hour. Then, the filters were manually removed before washing the individual pipette tips under high-pressure tap water. In step 2, the washed tips were sonicated at 40 kHz in an ultrasonic bath for 1 hour to remove small contaminants and particles from the surface of the pipette tip. In the final sterilization step, the tips were exposed to UV-C overnight or for 1 hour to remove infectious contaminants.

### Washing protocol 2. Pipette tip washing based on autoclave sterilization

Washing protocol 2 used in this study was aimed to mimic a generic protocol practiced in microbiology labs to reuse bio-contaminated glassware. The goal was to present an alternative washing protocol that individual labs can incorporate without reliance on a centralized washing station. Glassware with liquid culture is first autoclaved at 121 °C and 100 kPa with a 30-minute sterilization cycle and then bleached with 10% concentrated bleach. Next, the glassware is washed in Alconox detergent and then sterilized following two additional autoclave cycles. The first step is a liquid cycle of 30 min, followed by another round of dry cycle of 30 min. We followed similar washing and sterilization steps with slight modifications in our protocol to wash plastic pipette tips. Bleach and Alconox can deteriorate plastics at autoclave temperatures. To avoid plastic deterioration, we continued without bleach treatment and used high-quality dishwashing detergent to wash the pipette tips individually, as described in step 1 of washing protocol 1. Then, we loaded the washed tips in a reused pipette tip holder and autoclaved them for a 30-minute dry sterilization cycle.

### Extraction of T7 bacteriophage from pipette tips

To study the efficiency of the washing steps studied here, we extracted the residual phage from either unwashed or washed pipette tips by incubating them in the T7 phage extraction buffer. The T7 phage extraction buffer was prepared with 20 mM Tris-HCl, 100 mM NaCl, and 6 mM MgSO_4_ and then maintained at pH 8. We incubated the pipette tips from both washed and unwashed groups in phage extraction buffer from 2 hours to overnight at 4 °C.

### Quantification of T7 bacteriophage using standard double-layer plaque assay

We quantified the T7 phage titers extracted from the washed and unwashed pipette tips using the standard double-layer plaque assay (T7Select® System Manual, Novagen). A single colony of BL21 bacteria was grown in 4 ml M9LB agar media overnight at 37°C while shaking at 250 rpm. Then, we inoculated the fresh overnight culture in M9LB medium at a dilution of 1:100 and continued the shaking until the optical density (OD_600_) reached 1.0. A sufficient volume of top agarose was melted and maintained at 55°C. Simultaneously, a desired number of LB agarose plates were warmed to 37°C. Phage samples to be quantified from the ‘unwashed groups’ were diluted using ten serial dilutions in 90 µL of LB medium. Then, 200 µL of host cells were infected with 10 µL of each phage dilution. Since the phage titer in the ‘washed groups’ was expected to be low, 10 µL to 200 µL of undiluted phage samples were used to infect 200 µL of host bacterial culture to maximize the probability of plaque visualization. The infection was stopped within 5 minutes by adding 1.5 ml top agarose to the infection mixture before pouring the contents onto a prewarmed LB agarose 6-well plate. The poured plates were incubated for 3-4 hours at 37°C or overnight at room temperature for plaque formation. The plaques were counted manually, and the phage titer was calculated based on the dilution factor.

## RESULTS & DISCUSSION

TipNovus offers a high throughput and an automatic, continuous, UV-based washing and sterilization system for pipette tips. Although TipNovus is an efficient pipette tip decontamination option, it has yet to be accepted and implemented by research laboratories. One of the potential reasons is the high equipment cost, priced at $70,000, which can be unaffordable for individual academic research labs. Therefore, it may be beneficial to establish university-wide central pipette tip washing stations where biologically contaminated pipette tips can be sterilized. These washed tips can be reused in applications where an extremely sterile environment is not required or recycled appropriately. To advocate for a university-wide washing system, using UT Austin as an example university, we performed a cost-benefit analysis to reflect the economic, environmental, and energetic outcomes of installing the decontamination system (find detailed calculation in the **SI**).

Around 200 wet research laboratories at UT Austin use single-use plastic pipette tips. All the biologically contaminated pipette tips are collected for safe disposal by the campus Environmental Health and Safety (EHS). According to EHS, UT generates approximately 80,000 pounds of biologically hazardous waste annually. The collected biological waste is autoclaved by an external vendor and disposed of in a landfill in Buda, Texas. If the EHS were to recycle this waste, it would be comparable to saving 9,639 gallons of gasoline and 85.7 metric tons of carbon dioxide equivalent per year.

The 200 wet labs at UT spend more than $3 million a year to purchase plastic micropipette tips. Installing one TipNovus system would enable recycling 7% of UT’s pipette waste, saving $252,160 and resulting in a 100% return on investment within 70 days (**Figure 1a**). According to Grenova, the washed tips can be reused 5-40 times depending on the assay or the washing protocol used for the cleaning, therefore generating average cost savings of $112-$168 per hour.^12^ Based on this estimate, a conservative break-even period would be 625 hours of operation. TipNovus uses a pneumatic air compressor system, thus using limited electrical energy. According to our email conversation with Grenova, the equipment consumes electrical power at an amount equivalent to a coffee maker.^13^ If UT would operate the TipNovus equipment for 8 hours a day, then the electricity consumption to run the equipment would be 2920 kWh, resulting in an annual operating energy cost of $178. Similarly, the standard TipNovus machine consumes 1.5 gallons of water per washing cycle, resulting in an average water cost of $175 per year. Notably, the energy and water costs to manufacture a similar number of pipette tips are higher than TipNovus’s operating energy and water costs by 74 times and 37 times, respectively (**Figure 1b and 1c**). Therefore, the combined electricity and water cost to operate the washing system is estimated at $353 per annum, which is negligible compared to the pipette tip production cost.

**Figure 1.**
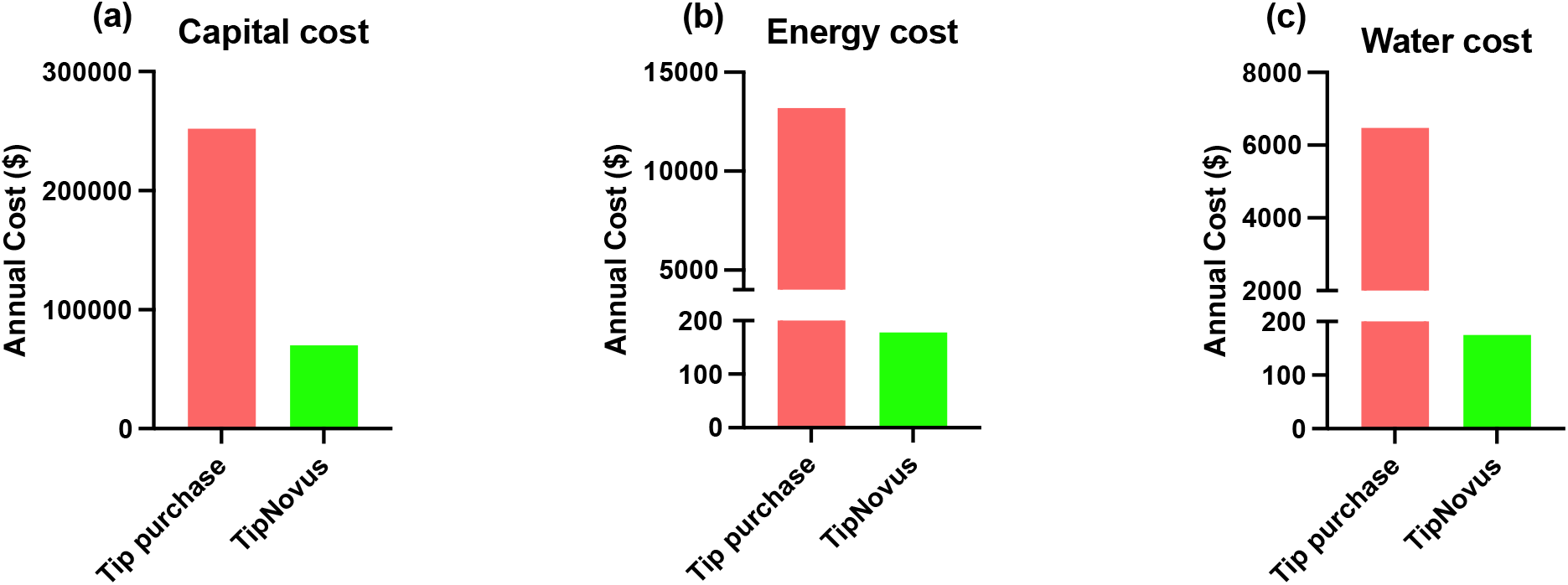
TipNovus washing installation provides economic and energy benefits for the University of Texas at Austin. (**a**) A single TipNovus washing station costs $70,000. Installing a machine at UT Austin is projected to reduce annual spending on pipette tips by $252,160 from pipette tip purchasing. TipNovus requires an annual (**b**) energy cost of $178 and a (**c**) water cost of $175 to run the equipment. To produce similar pipette tips, the energy and water requirements are 74 times and 37 times, respectively.

Along with TipNovus providing an environmentally sustainable solution to reduce plastic waste, it provides an economically and energetically viable solution. Because of UV-based sterilization, the water used for washing is already treated with UV, so the water waste can be drained directly or reused, limiting the generation of additional waste products. In contrast, wastewater disposal from plastic production remains a crucial environmental concern. Since the equipment installation cost can be returned from the one-time reuse of the washed and sterilized pipette tips, universities can take the initiative to install multiple university-wide washing stations to collect pipette tip wastes generated throughout the university and decontaminate them at a central facility. The university must employ staff to operate the equipment, which will incur additional running costs. Since the pipette tips can be reused several times, the resulting economic savings should make staff hiring feasible.

Another reason scientists are hesitant to accept the equipment is their reservations regarding reusing pipette tips after washing. From our discussion with many researchers, they have expressed that they are uncomfortable with reusing washed pipette tips for their sensitive experiments. The pipette tips washed by the TipNovus machine are being reused by many organizations for highly sensitive experiments, such as ELISA, NGS, PCR, mass spectroscopy, etc. The validation data comparing the results from fresh versus reused pipette tips is available upon request from Grenova. However, any direct evidence validating the efficiency of the washing system is currently unavailable on their website. Direct evidence of the effectiveness of this washing system will help convince universities’ EHS or sustainability departments to establish these university-wide washing stations. To perform an independent validation of the TipNovus washing protocol, we designed and performed a pilot study to directly estimate the sterilization efficiency of the washing system. We used T7 bacteriophage as a model contaminant for this pilot validation study. T7 bacteriophage or “phage” is a bacterial virus used as a platform for phage display technology. T7 phage particles are exceptionally robust and can survive in harsh conditions (temperature, pH, and chemicals) that inactivate other types of bacteriophages. They are highly resistant to storage and are shown to have retained viability even after 10-12 years at 4 ºC.^14^ They are most stable at a pH range from 6 to 8 for long-term storage but remain infective from pH 4 to 9 at least for 2 weeks.^15^ Moreover, the general phage quantification method, double layer plaque assay, utilizes significant micropipette tips. Therefore, we decided to use T7 bacteriophage as the biological contaminant in our pilot study. We examined whether implementing the TipNovus washing steps in series can eliminate phage residues from the contaminated tips.

We implemented the Grenova washing steps in series for our pilot study (see washing protocol 1 in materials & methods and **Figure 2a**). We estimated 0.07 ± 0.04% of the original phage stock retained per unwashed tip upon contamination. The washing steps with overnight UV sterilization removed 100% of the phage from the pipette tip surface (**Figure 2b**). Remarkably, we observed 100% phage removal from the pipette tips even when we reduced the UV-C sterilization time to 1 hour and doubled the initial phage stock (i.e., 4.80 × 10^9^ pfu) for contamination (**Figure 2c**). After washing the tips, followed by 1 hour of UV-C sterilization, we reused the washed tips to perform a standard double-layer plaque assay. The phage titer quantified with the reused tips was comparable to that estimated using fresh pipette tips (**Figure 2d**), confirming the washed tips’ reusability. The reusability of the tips further validates that the washing method is efficient. Although we did not estimate the residual phage DNA on the pipette tips, our study demonstrates the functional inactivity of the bacteriophage. Therefore, we reused the washed tips in an assay that relies on the functional activity of the phage.

**Figure 2.**
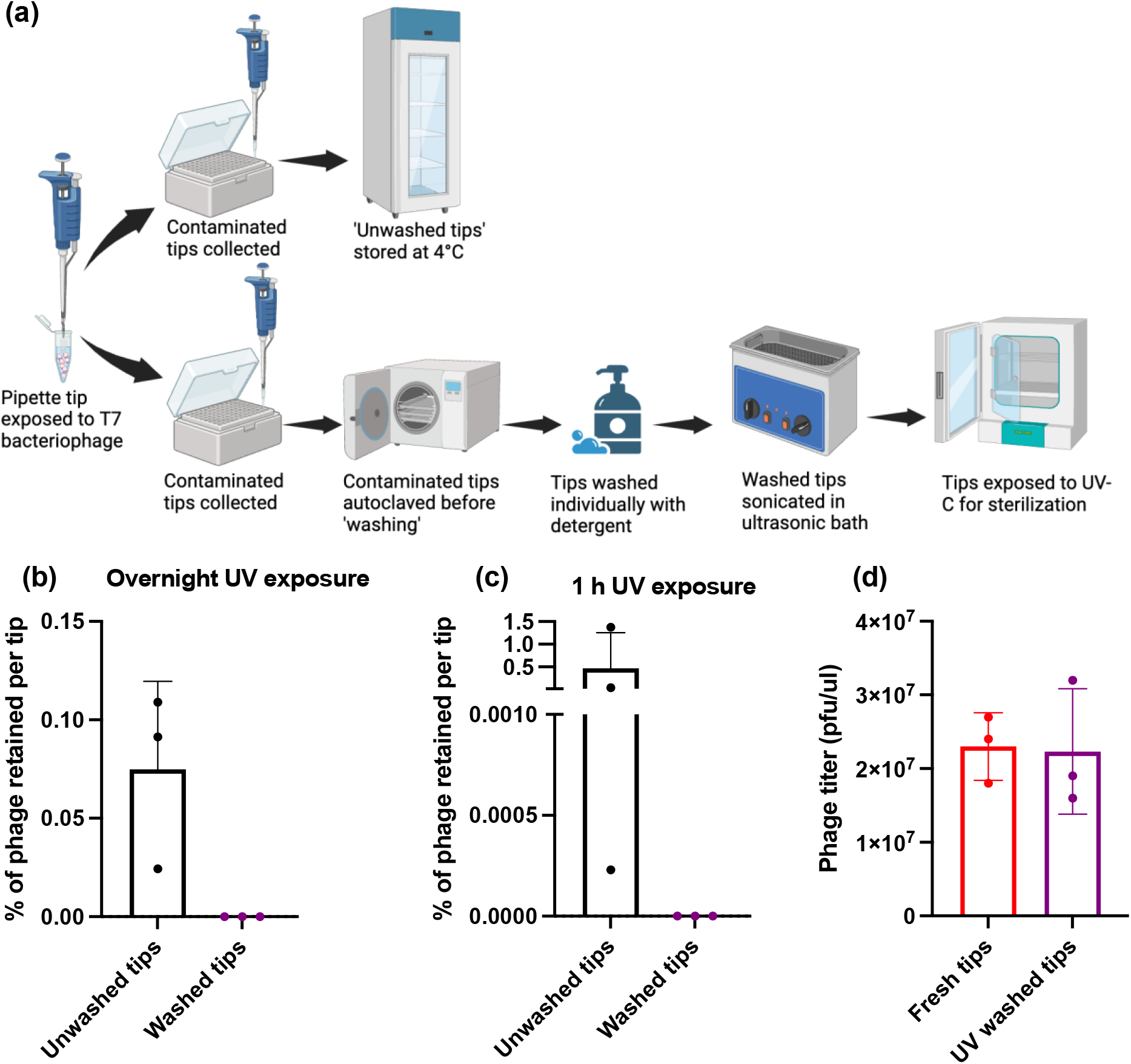
Washing and sterilization protocol following similar steps as TipNovus. (**a**) Washing protocol 1-We exposed the pipette tips to the T7 bacteriophage stock and collected them in equal numbers in two separate pipette tip racks. We stored one of the pipette tip racks at 4 °C until quantification and called them ‘unwashed tips.’ The other tip rack was first autoclave sterilized, washed individually, sonicated, and finally UV-sterilized, termed ‘washed tips.’ (**b**) Sterilizing pipette tips with overnight UV-C exposure efficiently removed 100% of the T7 bacteriophage from the tips, whereas the unwashed tips retained 0.07 ± 0.04% of phage stock per pipette tip. (**c**) Reducing the UV-C exposure to 1 hour also efficiently removed the T7 bacteriophage from the pipette tips. In contrast, the unwashed tips retained 0.47 ± 0.78% phage per pipette tip. (**d**) We identified an identical phage titer when the washed tips were reused for phage titer quantification as with fresh tips. Data represent mean ± SD (N=3). Two independent experimental repeats produced the same results.

We introduced an additional autoclave cycle before washing and sterilization steps (**Figure 2a**), which is not included in the TipNovus washing. We needed to include this step to handwash the tips after bacteria and phage exposure. The autoclave step has an additional effect on the pipette tip decontamination. Also, since we hand-washed the pipette tips individually, we manually removed the filters from the pipette tips. However, the high-pressure automatic TipNovus washing is not equipped for pipette tips with the filters. Bioresearch labs use many pipette tips without filters, which can be washed and sterilized efficiently with the TipNovus system.

As an alternative, we introduced and followed washing protocol 2 (**Figure 3a)**, an autoclave-based decontamination method. Here, we modified our washing steps to reflect cleaning practices that research labs routinely use to wash bio-contaminated glassware. We found that washing protocol 2 efficiently removed 100% phage from the contaminated tips, while the unwashed tips retained 0.47±0.78% of phage stock (**Figure 3b**). Then, we reused the autoclave-washed tips to estimate phage titer using the double-layer plaque assay and found the phage titers comparable to that estimated by fresh tips (**Figure 3c**). Therefore, the already established washing and sterilization protocol to reuse laboratory glassware can effectively wash bio-contaminated plastic pipette tips and potentially be reused.

**Figure 3.**
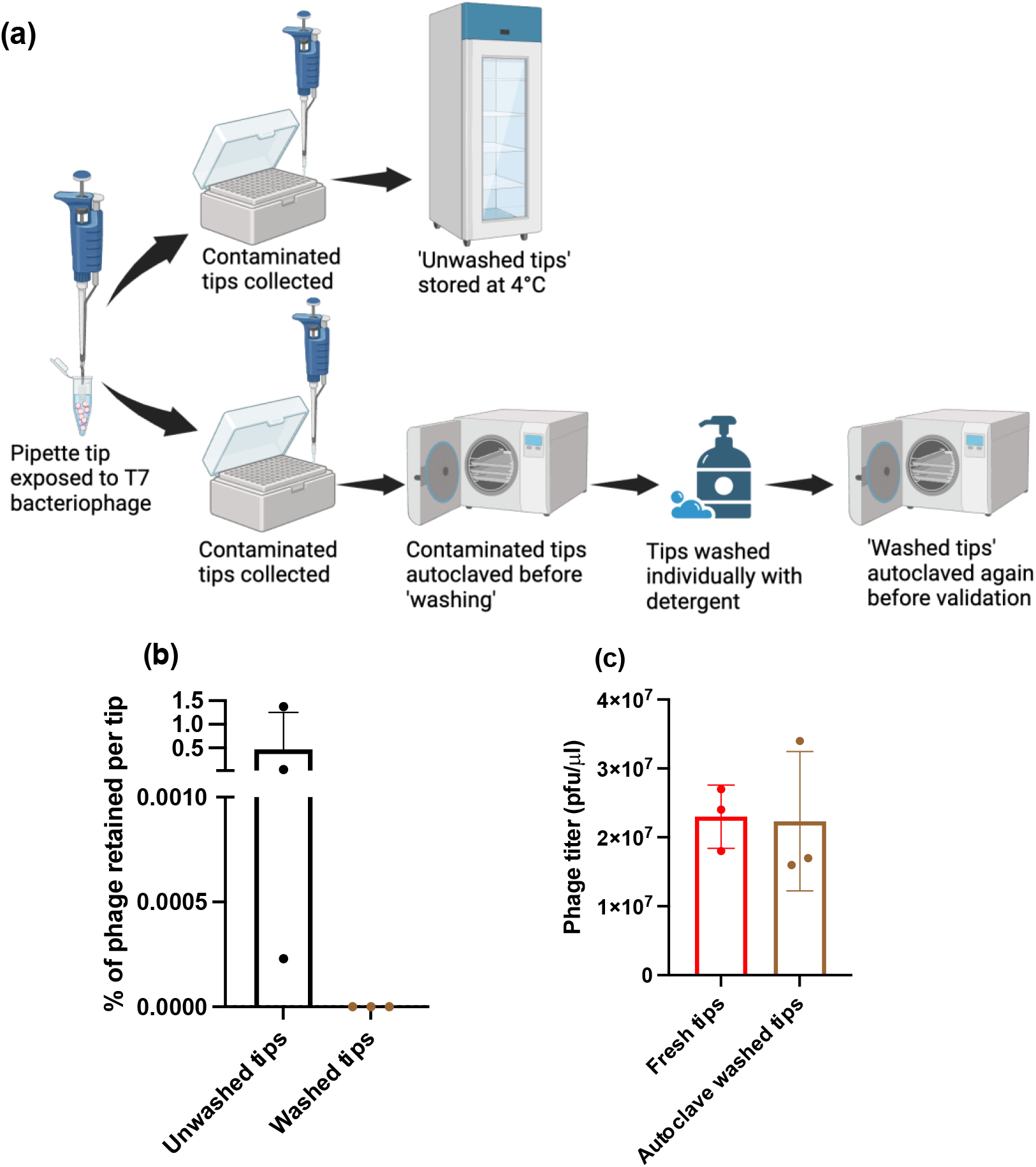
Washing and sterilization protocol following similar steps used to reuse glassware. **(a)** Washing protocol 2-We exposed the pipette tips to the T7 bacteriophage stock and collected them in equal numbers in two separate pipette tip racks. We stored one of the pipette tip racks at 4 °C until quantification and called them ‘unwashed tips.’ The other tip rack was first sterilized, washed individually, and autoclaved again after cleaning, termed ‘washed tips.’ (**b**) Decontaminating pipette tips following autoclave sterilization efficiently removed the T7 bacteriophage from the pipette tips, whereas the unwashed tips retained 0.47 ± 0.78% of phage stock per pipette tip. (**c**) Reusing washed tips estimated identical T7 phage titer in a stock solution compared to fresh tips using standard double-layer plaque assay. Data represent mean ± SD (N=3). Two independent experimental repeats produced the same results.

Since autoclave sterilization is already in place to decontaminate and reuse glassware in biological research labs, it can be conveniently implemented to wash pipette tips. Autoclave is one of the essential pieces of equipment for bioscientific researchers; thus, research laboratories or organizations already have it installed. However, the challenge with this washing strategy is that although the tip boxes can be sterilized in bulk, the tips must be washed individually in the current setting. It will be challenging to extrapolate the washing step to the tips waste generated from an entire lab or university. Remarkably, scientific laboratories from the developing world have created and implemented sustainable pipette tip-washing methods to address resource limitations while reducing plastic waste generation. A Bolivian researcher, Nataniel Mamani, has made an inexpensive tip washer from a plastic jar to reuse the pipette tips.^16^ The jar is designed with inner tubing that precisely fits the tips, enabling a high-pressure flow of water to wash out bleach and soap for thoroughly cleaning the tips. Similar high throughput pipette tip washing techniques can be designed and implemented to wash the contaminated tips in bulk. These combined approaches will not only lead to a substantial reduction in plastic waste but can also solve pipette tip requirements in resource-limited situations, such as the global tip shortage during the COVID-19 pandemic.

The current practice is to discard bio-contaminated pipette tips directly in the biohazard waste bin. For decontamination, scientists must eject them into an empty pipette rack. The contaminated tips must be segregated based on the contaminant and appropriately labeled. Then, the racks go into the TipNovus stage or autoclave for sterilization. This is a significant behavioral shift in the research culture as it can inconvenience scientists and affect experimental time points, leading to researchers hesitating about adopting the change. The university’s EHS and sustainability departments can help with initiatives to make the transition smooth and efficient. These initiatives will encourage scientists to practice sustainability in their research as passionately as they implement it in their personal lives.

This study encourages more research to verify the decontamination efficiency of washing stations for other biological contaminants. The success of these decontamination stations for plastic pipette tips could be extrapolated to other lab plastic wastes, such as falcon tubes, dishes, and plates. Although scientists have concerns about reusing sterilized plastics for sensitive experiments, these plastics can potentially be reused for less sensitive experiments (such as training) or recycled. More plastic reuse means less energy and water consumption, reduced plastic waste, and lower emissions from plastic distribution, limiting the overall environmental burden.

## Acknowledgment

This work is supported by Green Fund, a competitive grant program funded by UT Austin to support sustainability-related projects and initiatives proposed by students, faculty, or staff. We thank Dr. Debadyuti Ghosh for allowing us to use the Ghosh lab space to set up our experiments and for his guidance and support throughout the project. We acknowledge Jill Parrish, the Green Fund program coordinator, for coordinating the grant transitions and connecting us with the appropriate personnel to gather information for our cost-benefit estimation. Our special thanks to Patrick Mazur (senior facilities technical staff, UT Austin utilities, and energy management), UT Austin EHS department, and the representatives from Grenova for providing us with the information we used for the cost-benefit analysis. The schematics in the study are prepared using BioRender.

