## Supplemental Information for "Bio-contaminated Plastic Micropipette Tip Sterilization Stations: Environmentally, Economically, and Energetically Viable Solution"

**Table 1. Cost-benefit analysis of installing a TipNovus washing system at UT Austin**

| | |
|---|---|
| Overall plastic tip purchasing cost per year at UT | $3,631,104 |
| Estimated number of tip racks used per year for a small-sized wet lab | 2304 racks |
| Total number of wet labs in campus | 200 labs |
| Pipette tip cost per rack | $7.88 |
| (200uL non-filtered non-sterile tips from Thermofisher scientific for UT labs) | |
| Initial purchasing cost of a TipNovus equipment (Conversation with Grenova) | $70,000 |
| %age of tips possible to wash per year with one TipNovus machine at UT | 7% |
| Number of racks processed per cycle (From Grenova) (racks/cycle) | 4 racks |
| Processing time per cycle (From Grenova) (minutes/cycle) | 15 min |
| Number of racks washed per hour | 16 racks |
| Number of racks washed per day (Considering 8 hours running time per day) | 128 racks |
| Number of racks washed per year (Considering 250 working days per year) | 32,000 racks |
| Purchasing cost of 32000 pipette tip racks | $252,160 |
| Energy cost per year to run a TipNovus machine | $178 |
| Energy required to run TipNovus (if runs for 1 h per day) | 365 kWh/yr. |
| (According to Grenova, amount of energy used is comparable to a coffee maker) | |
| Energy cost of the equipment per year (8 hours running time per day) | 2,920 kWh |
| Electricity per kWh (Utilities and energy management department, UT Austin) | $0.061 |
| Energy cost to manufacture the same number of pipette tips | $13,197 |
| Energy consumption in producing plastic bottles (MJ/kg)[1] | 100 MJ/kg |

| | |
|---|---|
| Weight of one 1 L bottle (gm) | 38 g |
| Energy consumption to produce one bottle (MJ) | 3.8 MJ |
| Energy consumption to produce one tip (MJ) | 0.25 MJ |
| (Every 15 tips have same amount of plastic as in a bottle[2]) | |
| Number of tips washed per year (96 tips per rack) | 3,072,000 tips |
| Energy consumption to produce 3,072,000 tips (MJ) | 778,240 MJ |
| Energy consumption to produce 3,072,000 tips (kWh) | 216,350 kWh |

| Water cost per year to run a TipNovus machine | $175 |
|---|---|
| Water in gallons required for standard washing system per cycle (From Grenova) | 1.5 gallons |
| Estimated number of washing cycles to be required per year (4 cycles/hour) | 8,000 cycles |
| Water requirement per year (in gallons) | 12,000 gallons |
| Water cost per 1000 gallon | $14.55 |
| (Utilities and energy management department, UT Austin) | |

| Water cost manufacture the same number of pipette tips | $6,475 |
|---|---|
| Number of tips washed per year (96 tips per rack) | 3,072,000 tips |
| Number of bottle equivalent | 204,800 |
| (Every 15 tips have same amount of plastic as in a bottle[2]) | |
| Water use (It takes 8.23 liters to make one water bottle) (in liter) | 1,685,504 L |
| Water use during plastic tip preparation per year (in gallons) | 444,973 gal |

A TipNovus machine costs $70,000. The system is configured to work with manufactured tips from 10 μL up to 5 ml sizes. TipNovus can process four racks every 10-15 minutes, with a system throughput of 16-24 tip racks per hour. Conservatively, UT could process 32000 racks annually by installing one TipNovus machine with eight hours of operation, considering 250 working days a year. This would enable recycling 7% of UT's pipette waste and save $252,160. It will result in a 100% return on investment within 70 days, not accounting for the operating costs. TipNovus uses a pneumatic air compressor system, thus using limited electrical energy. According to our email conversation with Grenova, the equipment consumes electrical power at an amount equivalent to a coffee maker. An espresso coffee maker utilizes nearly 365 kWh of electricity annually when it runs for 1 hour per day.[3] If UT would operate the TipNovus equipment for 8 hours a day, then the electricity consumption to run the equipment would be 2920 kWh. The electricity cost per kWh is $0.061 (According to the Utilities and Energy Management Department, UT Austin), causing the annual operating energy cost to be $178. Similarly, the standard TipNovus machine consumes 1.5 gallons of water per washing cycle (information from Grenova). The water requirement for 2000 hours of operation would be 12000 gallons. Since 1000 gallons of water cost $14.55 (source- utilities and energy management department, UT Austin), an average water cost per year would be $175.

Notably, the energy and water costs to manufacture a similar number of pipette tips are higher than the operating energy and water cost of the equipment by 74 times and 37 times, respectively. We based our calculation on the total energy consumed in producing plastic bottles, which is about 100 MJ/kg when a 1L plastic bottle weighs 33g. To estimate the cost of water required for manufacturing, we used a reference that 15 pipette tips are equivalent to one plastic bottle based on the amount of plastic. Based on these assumptions, the energy cost to produce the pipette tips that one TipNovus can recycle over a year is $13,197, and the water cost is $6475. Therefore, the combined electricity and water costs to operate the washing system are estimated at $353 annually. This is negligible compared to the energy and water cost of producing the pipette tips.
